# Susceptibility of rosaceous fruits and apple cultivars to postharvest rot by Paecilomyces niveus

**DOI:** 10.1101/2021.04.01.438099

**Authors:** Tristan W. Wang, Kathie T. Hodge

## Abstract

Paecilomyces rot of apples is a postharvest disease caused by *Paecilomyces niveus*, a problematic spoiling agent of fruit juices and derivatives. The fungus produces ascospores that can survive food processing and germinate in finished fruit products. Processing apple fruits infected with Paecilomyces rot can lead to *P. niveus* contaminated juices. Because the fungus produces the mycotoxin patulin, juice spoilage by *P. niveus* is an important health hazard. Little is known about the disease biology and control mechanisms of this recently described postharvest disease. Following Koch’s postulates, we determined that a range of previously untested rosaceous fruits and popular apple cultivars are susceptible to Paecilomyces rot infection. We also observed that two closely related food spoiling fungi, *Paecilomyces fulvus* and *Paecilomyces variotti*, were unable to infect, cause symptoms in, or reproduce in wounded fruits. Our results highlight the unique abilities of *Paecilomyces niveus* to infect a variety of fruits, produce patulin, and form highly-resistant spores capable of spoiling normally shelf-stable products.

## Introduction

Patulin is a serious mycotoxin that can cause gastrointestinal, immunological, and neurological damage, and is an important contaminant of apple products and their derivatives (Puel et al. 2010). One 2009 survey detected patulin in 23% of apple juices and ciders sampled from Michigan retail grocery stores, with 11.3% of these samples containing patulin over the FDA limit of 50ppb (Harris et al. 2009). Contamination by this mycotoxin is thought to occur through postharvest fruit infection and food spoilage by patulin-producing microbes. At least 18 *Penicillium, Aspergillus*, and *Paecilomyces* species are known to produce patulin (Puel et al. 2010) and the mycotoxin has been detected in a wide variety of other fruit products including pear, peach, cherry, apricot, orange, and mango juices and jams (Erdogan et al. 2018, Hussain et al. 2020, Moake et al. 2005, Spadaro et al. 2008).

One important patulin producer, *Paecilomyces niveus* Stolk & Samson (*Byssochlamys nivea* Westling), uniquely threatens juice production as it is a thermotolerant mold that creates durable ascospores. The ascospores are the predominant structures responsible for *P. niveus* food contamination. Contamination by *P. niveus* is a long-standing issue and the fungus has been found to contaminate a wide variety of fruit products including apple-based products, concentrated orange juice, strawberry puree, and tomato paste (Kotzekidou 1997, dos Santos et al. 2018).

*Paecilomyces niveus* is a common soil fungus, found present in a third of New York orchard soil samples (Biango-Daniels and Hodge 2018). It has been thought that *P. niveus* contamination of food products is environmental, originating from soil, air, and equipment. Even small quantities of this homothallic fungus pose a hazard for juice production as a single *P. niveus* spore can give rise to heat-resistant ascospores. Recently, *P. niveus* was identified as the causal agent of Paecilomyces rot, a postharvest disease of apples (Biango-Daniels and Hodge 2018) and citrus fruits (Wang and Hodge 2020). Biango-Daniels et al. (2019) showed that processing apple fruits infected with Paecilomyces rot can result in apple juice significantly contaminated with patulin and viable *P. niveus* ascospores. Since the description of Paecilomyces rot (Biango-Daniels and Hodge 2018), *Paecilomyces niveus* has been found infecting apples in a fruit market in China (Khokhar et al. 2019). In addition, the fungus was observed growing on peas in Serbia (Dragic et al. 2016) and on aphids in Brazil (Zawadneak et al. 2015).

The broad range of fruit products in which *P. niveus* and patulin have been found led us to hypothesize that the fungus may be able to infect and reproduce in a range of fruits. In addition, we sought to further characterize the disease biology of Paecilomyces rot by testing the susceptibility of other popular apple cultivars (Empire, Fuji, Granny Smith, and Golden Delicious). Lastly, we asked whether pathogenesis in our wound challenges was unique to *P. niveus*. Like *P. niveus*, two closely related heat-resistant molds known to contaminate fruit products, *Paecilomyces fulvus* and *Paecilomyces variotii*, produce heat-resistant ascospores capable of surviving temperatures above 85°C (Houbraken et al. 2006, Samson et al. 2009). We hypothesized that both can also cause symptoms, reproduce in apple fruits, and contaminate apple products via infected apple fruits.

To test the preceding hypotheses, we tested the wound-infecting ability of *P. niveus* in fruits of a variety of apple relatives: peaches, pears, sweet cherries and sour cherries. In addition, we inoculated and compared lesion development in four popular apple cultivars. Lastly, to test for the disease-causing abilities of *P. niveus* relatives in wounded apple fruits, we inoculated apple fruits with *Paecilomyces variotii* and *Paecilomyces fulvus* and observed them for symptom development.

## Materials and methods

### Fruit

Detached peach fruits (*Prunus persica*, cv. Lori Anne) (n=26) and pear fruits (*Pyrus communis*, cv. Green D’Anjou) (n=31) were selected at a local supermarket based on their uniformity in size, and absence of both wounds and disease symptoms. Sour cherries (*Prunus cerasus*, cv. Montmorency) (n=40) and sweet cherries (*Prunus avium*, cv. Kristin) (n=31) were freshly picked from a local New York orchard. For apple cultivar susceptibility testing, Empire (n=31), Fuji (n=28), Golden Delicious (n=31), and Granny Smith (n=31) apples were purchased from a local supermarket. These four cultivars were chosen based on their popularity in US markets. Three of these cultivars (Empire, Fuji, and Granny Smith) have not previously been tested for susceptibility to Paecilomyces rot. For susceptibility tests involving *P. fulvus* and *P. variotii*, 30 Empire apples were used for inoculation of each fungus.

### Fungal pathogens

*Paecilomyces niveus* strain CO7, isolated from culled apple fruits in New York, was used to inoculate fruits using the method described in Biango-Daniels and Hodge (2018). We have sequenced the full genome of this strain (Biango-Daniels et al. 2018), and found that sequences from ITS and BenA regions can reliably be used to confirm identity as *P. niveus. P. niveus* strain MC4, isolated from New York residential garden soil and *P. niveus* strain 106-3, isolated from New York orchard soil, were also used in fruit inoculation and identified using their ITS and BenA regions. *Paecilomyces fulvus* strain 7, obtained from the Worobo lab collection and originally isolated from spoiled food and *Paecilomyces variotii* strain 103-2, isolated from NY soil, were identified morphologically and by sequencing of the ITS region.

To produce inoculum, fungi were grown from 5mm plugs taken from the edge of 2-week-old colonies on PDA. Five plugs of *P. niveus* were cultured on PDA. The fungus was allowed to grow in the dark for two weeks at 25°C, covering tyndallized toothpicks that had been autoclaved twice: once in water and once in potato dextrose broth the following day. Control toothpicks were similarly treated, but in the absence of the fungus. Both *P. fulvus* and *P. variotii* were grown as described above to produce toothpick inoculum.

### Fruit handling, inoculation, and measurement

Fruits were sanitized before fungal inoculation: Peach fruits were sterilized for 30s in 70% ethanol, 2 min in 1% sodium hypochlorite, and 15s in 70% ethanol and then left to dry in a Class IIB biological safety cabinet. Two treatment toothpicks and two control toothpicks were inserted on opposite sides of the fruit’s equator. The toothpicks served to both wound (1 cm deep) and inoculate the fruits. Individual fruits and wounds were numbered. The peaches were placed in moist chambers in a dark incubator at 25°C. Over the course of 2 to 3 weeks, fruits and disease symptoms were observed. Horizontal and vertical diameters of lesions were quantified using a digital caliper (VWR Carbon Fiber Composite) every other day. Fruits displaying colonization or symptoms such as blue-green or brown sporulation, indicative of other common postharvest diseases, were removed from the study and further analysis. Pear and apple fruits were similarly sanitized and inoculated. Individual fruits of the four apple cultivars were randomized in moist chambers. Both sour and sweet cherry fruits were wounded only two times on opposite sides (5 mm deep), once with a treatment toothpick and once with a control toothpick on opposite sides of the fruit, before being laid to rest in moist chambers. To test for fruit susceptibility to other *P. niveus* strains, *P. niveus* MC4 and 106-3 were each used to inoculate two Empire, Fuji, and Granny Smith apples in addition to three D’Anjou pears.

### Statistical analysis

To test for significance of lesion diameters, statistical analysis was done using two separate linear mixed-effects models at day 6 and day 20 for apple fruits, day 2 and at day 12 for pear fruits, and at day 2 and day 8 for peach and cherry fruits. These models were constructed in R statistical programming software using the lmerTest and emmeans packages. Ggpubr, and ggplot2 packages were used to construct models and visualize results. The models took into consideration the fixed effects of the incubation chamber and the random fruit identification number. To test for differences in cultivar resistance, differences in lesion growth were measured using a one-way ANOVA, followed by a post hoc Tukey HSD test.

## Results

### Lesion growth in peaches, pears, and cherries

In all fruits tested for wound infection with *P. niveus*, spreading lesions were observed and lesion diameters (± standard error) were measured every other day (Fig. 1). In peaches, pears, and cherries, lesions developed at every inoculation point and grew rapidly over the course of 2 weeks. The average lesion diameters in peaches, sweet cherries, and sour cherries at day 2 were compared to average lesion diameters at day 8 (P<2e-16). Pear lesions at day 2 and 12 were similarly compared (P<2e-16). At two days, average treatment lesion diameters were 3.76 ± 0.54 mm (n=26) for Lori Anne peaches, 2.93 ± 0.25 mm (n=31) in D’Anjou pears, and 13.32 ± 0.86 mm (n=40) in Montmorency cherries. No lesions were detected in Kristin cherries at two days. At 8 days postinoculation, average treatment lesion diameters were 30.01 ± 1.98 mm (n=14) for Lori Anne peaches, 16.51 ± 0.45 mm (n=29) for Montmorency cherries, and 13.32 ± 0.86 (n=18) for Kristin cherries. At 12 days, average treatment lesion diameter was 49.15 ± 0.41 mm (n=30) in D’Anjou pears.

**Fig. 1.**
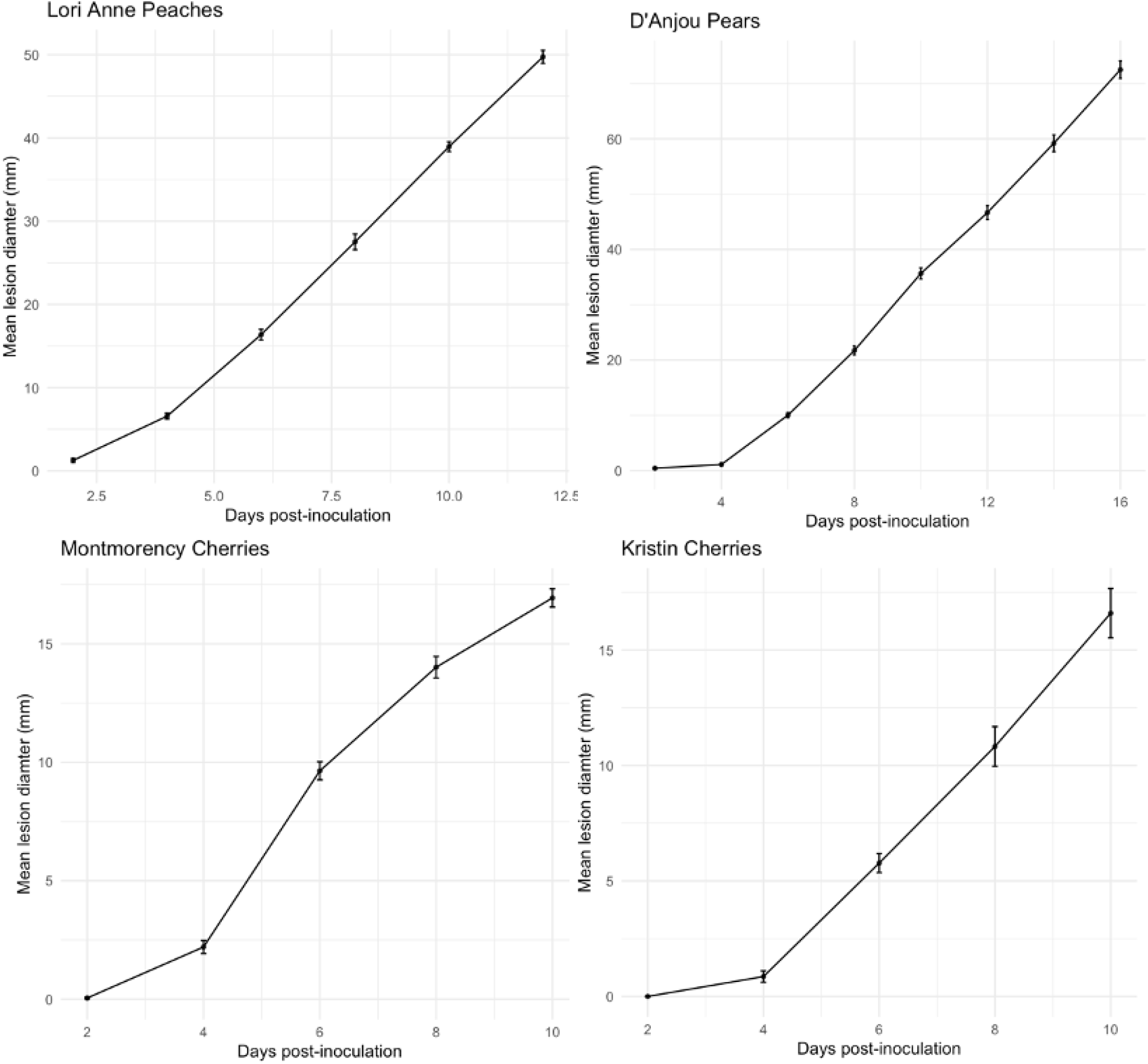
Measurements of mean lesion diameter (± 1 standard error) over the course of 2 to 3 weeks incubation (25°C, ≥95% humidity) on Lori Anne peaches, D’Anjou pears, Montmorency sour cherries, and Kristin sweet cherries inoculated with *Paecilomyces niveus*. Lesions on control toothpicks were nearly absent in peaches and pears and completely absent in both Kristin and Montmorency cherries.

Towards the end of the experiment, three out of 24 control toothpicks used in peaches on day 12 and one control toothpick out of 60 in pears on day 14 showed lesion development, likely due to infection by a different postharvest pathogen, and were taken out of the experiment and statistical analysis. No control toothpick in sweet and sour cherries showed detectable lesion development throughout the experiment. Several fruits, particularly Lori Anne peaches and Kristin sweet cherries, displayed symptoms of other postharvest infections in the duration of the study and were removed from the study and further analysis.

### Characterization of Paecilomyces rot in Rosaceae fruits

In Lori Anne peaches, circular lesions developed at the site of inoculation (Fig. 2A). After 7 days postinoculation, discoloration of the epidermis was light brown and the lesion was semi-firm to the touch. Unlike the distinct lesion borders found in Paecilomyces rot in apples, lesion borders in peaches were less distinct. White, branching surface hyphae could be seen in early stages of the disease (1 week postinoculation), and extended deeply into the fruit. Fruit flesh discoloration ranged from yellow-brown to brown. Necrotic tissue was not easily separable from healthy tissue.

**Fig. 2.**
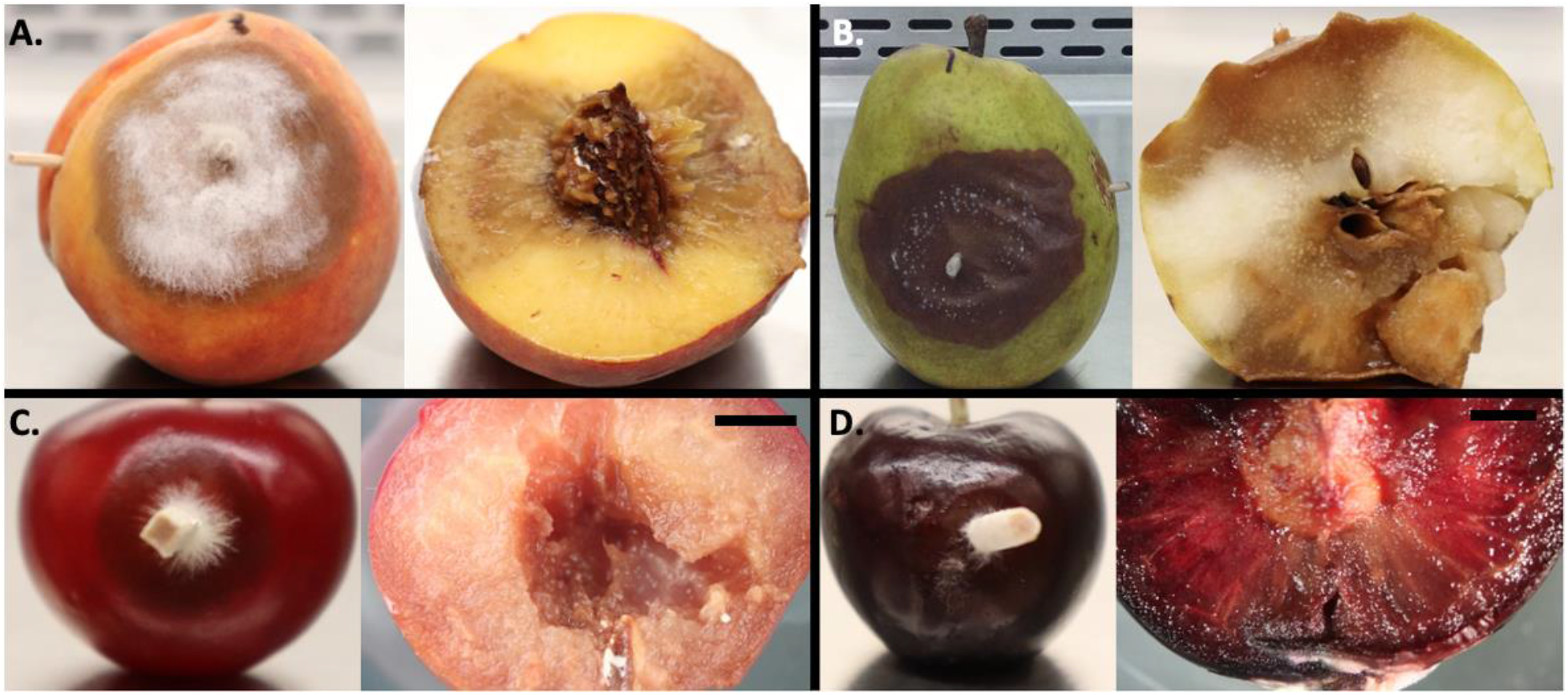
External and internal symptoms of infected rosaceous fruits. Infected fruits, two weeks postinoculation after incubation in dark, moist chambers (25°C, ≥95% humidity) of (**A**) Lori Anne peaches and (**B**) D’Anjou pear with cross-sections of each. Close-up view of external developing lesion and cross-section showing internal rot on (**C**) Montmorency and (**D**) Kristin cherry. Scale bars are 3 mm.

D’Anjou pear infections resulted in dark brown lesions that were generally circular and even in discoloration (Fig. 2B). Lesions with sharp, distinct edges expanded radially from the point of inoculation. Unlike Paecilomyces rot in apples which manifests as a hard rot and is spongy to the touch, necrotic tissue within pear fruits was soft, wet, and fragile. Fruit epidermis on the lesion surface became papery and brittle. After 14 days postinoculation, faint concentric rings of varying brown shades and tufts of white hyphae can be observed at lenticels. Yellow-brown internal rot spread deeply into the fruit. Diseased tissue was easily separable from healthy tissue. Unique to pears, diseased flesh became translucent and extremely soft, which allowed for clear visibility of pear sclereids.

In both Montmorency sour and Kristin sweet cherries (Fig. 2C and 2D), circular lesions developed at the site of inoculation. Epidermal discoloration was brown, sometimes light-brown close to the center of the lesion in sour cherry lesions and yellow-brown in sweet cherry lesions. While lesion edges are distinct in sour cherries, lesion edges were less pronounced in sweet cherries. After 14 days postinoculation, concentrated tufts of mycelia appeared on the surface of sweet cherries. Rot extended deeply into the fruit in both sweet and sour cherries resulting in an orange-brown to brown flesh discoloration. Necrotic tissue was soft and watery and not easily separated from healthy tissue.

### Completing Koch’s postulates

For peaches, pears, and cherries newly-tested for Paecilomyces rot, lesion surfaces of two individual symptomatic fruits were sterilized with 70% ethanol. Infected interior flesh was cultured for fungus. The isolated culture was then used to infect healthy fruits, and lesions developed that were identical to those previously observed in the source peach, pear, and cherry fruits. *P. niveus* was cultured from the newly infected fruits and identified morphologically by its naked asci and white colonies that yellow with age, thereby completing Koch’s postulates (Samson et al. 2009).

### Apple cultivar-based susceptibility and *P. niveus* lesion size

All four apple cultivars inoculated with *Paecilomyces niveus*, Golden Delicious, Empire, Granny Smith and Fuji (the latter three have not previously been tested for susceptibility to Paecilomyces rot), showed clear signs of lesion development 4 to 6 days postinoculation. Over the next two to three weeks, lesions continued to grow rapidly and average diameters of treatment lesions were measured every other day. At six days postinoculation, lesion diameters (± standard error) were 6.44 ± 0.68 mm (n=31) on Empire, 9.30 ± 0.61 mm (n=28) on Fuji, 6.86 ± 0.47 (n= 31) on Golden Delicious, and 10 ± 0.73 (n = 29) on Granny Smith apples (Fig. 3A). At 20 days postinoculation, lesion diameters were 51.98 ± 4.28 mm (n = 31) on Empire, 33.52 ± 1.62 mm (n = 26) on Fuji, 14.49 ± 1.36 (n= 31) on Golden Delicious, and 33.38 ± 2.57 (n = 29) on Granny Smith apples (Fig. 3A and 3B).

**Fig. 3.**
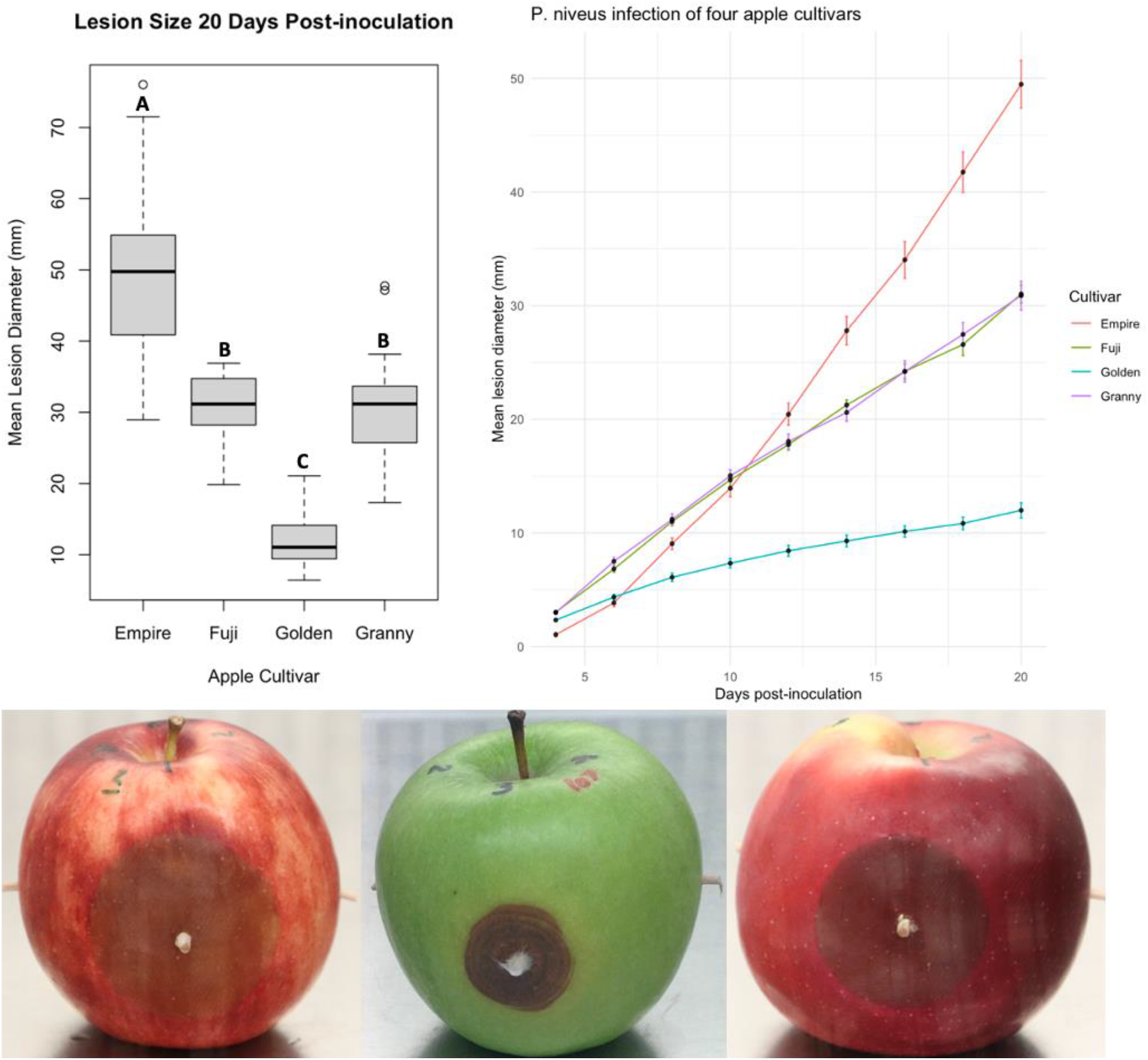
Comparison of Paecilomyces rot infected apple cultivars. (a.) 16-day progression of mean lesion diameter of four apple cultivars stored in the dark (25°C, ≥95% humidity) infected with *P. niveus*. (b.) Mean lesion diameters on infected apple cultivars at 20 days postinoculation. (c.) From left to right: infected Empire, Granny Smith, and Fuji apples after two weeks postinoculation.

### Pathogenicity of *Paecilomyces fulvus* and *P. variotii* in apple fruits

We investigated whether two closely related *Paecilomyces* species, *P. fulvus* and *P. variotii*, can also infect and grow in apple fruits. After three weeks, no noticeable symptoms developed in Empire apples inoculated with either fungus. Apple fruits remained intact, and at the end of the three-week period were cut in half for further observation. Wounding created by the toothpick insertion remained identical to the control toothpick wound, and no rot or lesions developed. There was no evidence these fungi were able to infect the apple.

## Discussion

The Rosaceae is a large plant family that includes economically important fruit crops such as peaches, pears, cherries, strawberries, and apples, the latter being the most consumed fruit in the United States (USapple). Each year the US produces 240 million bushels of apples, a third of which are processed into juices and other apple derivatives (USapple). In 2020/2021, the United States also produced 658,000 tons of pears, 720,000 tons of fresh peach and nectarines, and 383,000 tons of cherries (USDA-FAS 2020a,b). Postharvest diseases are responsible for a 20– 25% loss of harvested fruits and vegetables in developed countries and even more in developing countries (Nunes 2012). A 2018 study reported that 92% of juice manufacturers surveyed have experienced fungal spoilage in finished products (Snyder and Worobo 2018). *Paecilomyces* is a notorious genus that includes both important food spoiling fungi (*P. niveus, P. fulvus, P. variotii*) and emerging opportunistic pathogens of humans (*P. variotii* and *P. formosus*) (Heshmatnia et al. 2017). Unlike common postharvest pathogens of apples, *Paecilomyces niveus* uniquely threatens juice production because it can not only produce the FDA-regulated mycotoxin patulin, but can also infect apples as a postharvest disease and persist through juice sterilization processing. Spoilage by *P. niveus* directly leads to contamination of finished food products with patulin, and potentially other mycotoxins including byssochlamic acid, byssochlamysol, and mycophenolic acid (Houbraken et al. 2006).

Data presented in this study suggest that *P. niveus*, a patulin-producing, heat-resistant mold, can also be a wound-infecting pathogen of rosaceous fruits other than apples. *Paecilomyces niveus* strain CO7, isolated from decaying apples, was able to grow, reproduce, and cause symptoms in other fruits including pears, peaches, and both sweet and sour cherries. Two additional *P. niveus* strains, MC4 and 106-3, were also confirmed through Koch’s postulates to be able to infect and reproduce in Empire, Fuji, Granny Smith apples and D’Anjou pears. We also generated preliminary data to suggest that *P. niveus* can also rapidly infect strawberries and raspberries, but these infections were often masked by other postharvest diseases, complicating analysis. Paecilomyces rot manifested similarly across all these fruits, causing lesions and internal rot at wound sites, consistent with the original description of the disease (Biango-Daniels and Hodge 2018). Key differences in symptom development included profuse mycelial growth on the surface of peaches, dense tufts of mycelia in cherries, and notable disintegration of fruit flesh in pears. We observed that fruit disintegration from *P. niveus* infection occurred at a faster rate in peach and pear fruits than it did in apple fruits. In addition, all apple cultivars inoculated with *P. niveus*, were susceptible to Paecilomyces rot infection. Interestingly, the rate of lesion growth in inoculated Golden Delicious apples was slower than previously observed (Biango-Daniels and Hodge 2018) suggesting that other external factors may influence lesion development.

We also tested the infectious abilities of two close relatives of *Paecilomyces niveus: Paecilomyces fulvus* and *Paecilomyces variotii*. Neither fungus was observed to cause disease, despite their close relationship with *P. niveus*, and their status as common heat-resistant spoilage fungi of processed fruit products. These data provide a contrast to the infectious abilities of *P. niveus*, a pathogen that can grow and reproduce in living fruits.

Our findings have implications for fruit growers and juice producers as they reveal that diseased peaches, pears, and cherries can harbor large amounts of *P. niveus* spoilage inoculum. And since *P. niveus* ascospores tolerate high heat, they can survive thermal processes, especially when suspended in fruit products like strawberry puree (Silva 2015), pineapple juice (Ferreira et al. 2009), canned tomato paste (Kotzekidou 1997), apple, and cranberry juice (Palou et al. 1998).

This study disproves the prevailing belief that *P. niveus* contamination can occur solely from environmental sources, and suggests that diseased fruits can be a source of spoilage inoculum. The sporadic nature of food spoilage makes it difficult to demonstrate this route of contamination in real industrial processes. Our results also broaden the known host range of *P. niveus* to include economically important and commonly processed fruits including peaches, pears, sweet cherries, and sour cherries, and show that multiple popular apple cultivars are susceptible to infection. Future work should address detection of infected fruits in the field and the risk that *P. niveus* infection of fruits can be a source of spoilage inoculum.

## Acknowledgements

We would like to thank Dr. Erika Mudrak for her assistance with statistical analyses. Aarzoo Haideri, Amanda Wilson, and Eshan Mehrotra assisted with fruit inoculation, lesion measurements, and photographs. Dr. Megan Biango-Daniels and the Worobo lab generously provided *Paecilomyces* cultures.

